# Abundant expression of maternal siRNAs is a conserved feature of seed development

**DOI:** 10.1101/866806

**Authors:** Jeffrey W. Grover, Diane Burgess, Timmy Kendall, Abdul Baten, Suresh Pokhrel, Graham J. King, Blake C. Meyers, Michael Freeling, Rebecca A. Mosher

**Affiliations:** Department of Molecular and Cellular Biology, The University of Arizona, Tucson, AZ, 85721 USA; Department of Plant and Microbial Biology, The University of California, Berkeley, CA, 94720 USA; The School of Plant Sciences, The University of Arizona, Tucson, AZ, 85721 USA; Southern Cross Plant Science, Southern Cross University, Lismore, NSW, 2480 AUS; AgResearch Ltd, Grasslands Research Centre, Palmerston North, New Zealand; Donald Danforth Plant Science Center, St. Louis, MO 63132 USA; Division of Plant Sciences, University of Missouri – Columbia, Columbia, MO 65211 USA; Bio5 Institute, The University of Arizona, Tucson, AZ, 85721 USA

## Abstract

Small RNAs are abundant in plant reproductive tissues, especially 24-nt short interfering (si)RNAs. Most 24-nt siRNAs are dependent on RNA Pol IV and RDR2, and establish DNA methylation at thousands of genomic loci in a process called RNA-directed DNA methylation (RdDM). In *Brassica rapa*, RdDM is required in the maternal sporophyte for successful seed development. Here we demonstrate that a small number of siRNA loci account for over 90% of siRNA expression during *B. rapa* seed development. These loci exhibit unique characteristics with regard to their copy number and association with genomic features, but they resemble canonical 24-nt siRNA loci in their dependence on RNA Pol IV/RDR2 and role in RdDM. These loci are expressed in ovules before fertilization and in the seed coat, embryo, and endosperm following fertilization. We observed a similar pattern of 24-nt siRNA expression in diverse angiosperms despite rapid sequence evolution at siren loci. In the endosperm, siren siRNAs show a marked maternal bias, and siren expression in maternal sporophytic tissues is required for siren siRNA accumulation. Together these results demonstrate that seed development occurs under the influence of abundant maternal siRNAs that might be transported to, and function in, filial tissues.

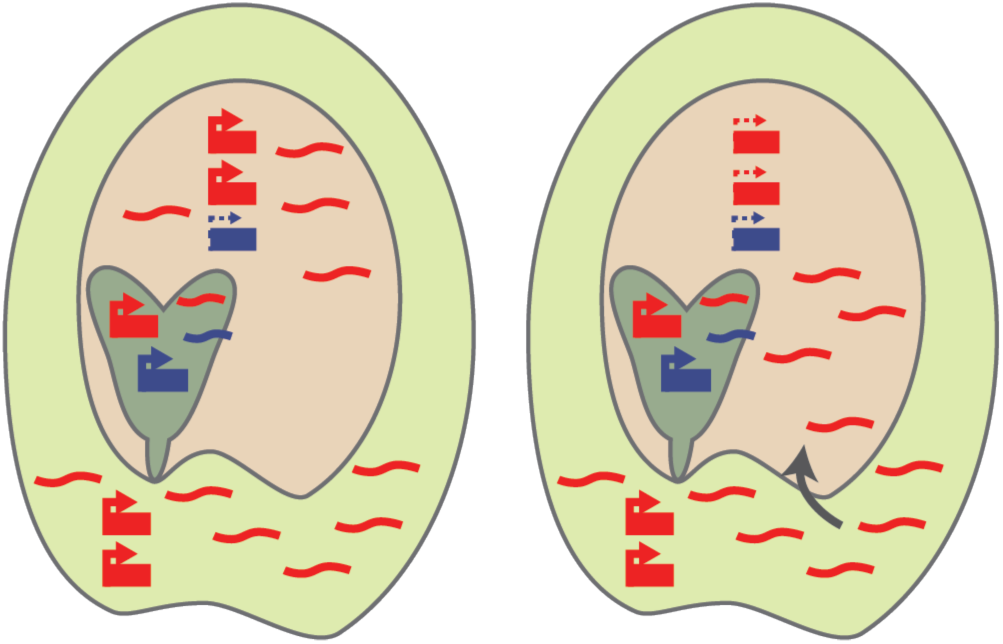

## Introduction

Seeds are critical for the worldwide food supply. Therefore, factors which influence seed development underpin global food security and represent important avenues for crop improvement. Seed development is complex, as seeds are composed of multiple genetically-distinct tissues, including the embryo and endosperm, which are products of fertilization of the haploid egg and diploid central cell, respectively. The seed coat surrounds and protects the embryo and endosperm, is entirely maternal and sporophytic in origin, and is descended from the integument tissue surrounding the female gametes. Several studies indicate that 24-nt small interfering (si)RNAs and RNA-directed DNA Methylation (RdDM) are important during early seed development, although a biological role for RdDM during seed development has remained elusive (Erdmann et al., 2017; Kirkbride et al., 2019; Lu et al., 2012; Martinez et al., 2018; Satyaki and Gehring, 2019).

A hallmark of RdDM is accumulation of 24-nt siRNAs produced by RNA Pol IV and RNA-DEPENDENT RNA POLYMERASE 2 (RDR2). RNA Pol IV and RDR2 together produce short double-stranded transcripts (Blevins et al., 2015; Zhai et al., 2015). Pol IV/RDR2 products are then cleaved by DICER-LIKE 3 (DCL3) to produce 24-nt siRNA duplexes (Singh et al., 2019) that can be bound by ARGONAUTE 4 (AGO4), which removes the “passenger strand,” leaving the “guide strand” bound (Havecker et al., 2010; Wang and Axtell, 2017). The AGO4-siRNA complex interacts with the C-terminal domain of RNA Pol V and either the Pol V transcript or DNA within the Pol V transcription bubble (El-shami et al., 2007; Lahmy et al., 2016; Li et al., 2006; Liu et al., 2018), leading to the recruitment of the DNA methyltransferase DRM2 and methylation of cytosines (Matzke and Mosher, 2014). RdDM activity is associated with transcriptional silencing of transposable elements in euchromatin (Stroud et al., 2013; Zemach et al., 2013), and can influence the expression of genes (Grover et al., 2018; Pontier et al., 2005).

Despite the abundance of 24-nt siRNAs in seeds, seed development proceeds normally in *Arabidopsis thaliana* lacking 24-nt siRNAs or RdDM machinery. However, loss of RdDM increases viability in paternal-excess crosses, suggesting that balancing the parental contributions of siRNAs might be important for seed development (Erdmann et al., 2017; Martinez et al., 2018; Satyaki and Gehring, 2019). In *Brassica rapa*, a close relative of *A. thaliana*, RdDM mutants display a single and specific defect – seed abortion approximately 15 days post fertilization (Grover et al., 2018). This phenotype is controlled by maternal sporophyte genotype, further supporting the hypothesis that parental siRNA contributions are important for seed development.

Previous work has identified both maternal- and paternal-specific siRNAs in reproductive tissues. Male gametophytes (pollen grains) produce Pol IV-dependent, epigenetically activated small interfering RNAs (easiRNAs) that are proposed to enter filial tissues during fertilization (Martinez et al., 2018; Martínez et al., 2016; Slotkin et al., 2009). However, the maternal genome is the source for the majority of 24-nt siRNAs in the developing seed, including a class of siRNAs which exclusively accumulate in reproductive tissues (Mosher et al., 2009; Pignatta et al., 2014). Maternal siRNA accumulation in the seed is linked to expression of developmental regulators in *A. thaliana* endosperm (Kirkbride et al., 2019; Lu et al., 2012). It has been hypothesized that siRNAs are imprinted in the endosperm (Lu et al., 2012; Mosher et al., 2009; Rodrigues et al., 2013), though perturbation of known imprinting pathways did not change their apparent uniparental expression pattern (Mosher et al., 2011), and this interpretation has been challenged (Pignatta et al., 2014). Rather than imprinted expression, maternal siRNAs could be loaded into developing filial tissues from the surrounding maternal sporophytic tissue, in a manner similar to maternal loading of piRNAs into *Drosophila melanogaster* eggs before fertilization (Brennecke et al., 2008; Klattenhoff and Theurkauf, 2007). However, such a mechanism has yet to be observed in plants.

Here, we demonstrate that the small RNA content of developing seeds is predominantly comprised of a distinct category of maternally-expressed siRNAs. These siRNAs accumulate at a small number of genomic loci we call “sirens,” based on a category of small RNAs with characteristics similar to our own (Rodrigues et al., 2013). Siren siRNAs are highest expressed in the ovule before fertilization and the developing seed coat after fertilization, and might move between maternal and filial tissues. Siren siRNAs trigger DNA methylation, uncovering a potential mechanism for maternal sporophytic control over seed development. Furthermore, we find siren loci in multiple species and rare conservation of siren position, but not sequence, among crucifers, indicating that siren loci are a general feature of angiosperm seed development. These results resolve conflicting reports regarding the origin of siRNAs in the seed and illuminate mechanisms for parental conflict and transgenerational inheritance.

## Results

### *B. rapa* ovule siRNAs overwhelmingly accumulate within “siren” loci

To better understand the role of siRNAs during seed development, we sequenced small RNAs from a variety of *B. rapa* tissues, organs, and organ systems: whole seed samples collected throughout development (10, 14, 19, and 24 days after fertilization), the constituent parts of the seed (embryo, endosperm, and seed coat), and mature leaves. Together with our previously published *B. rapa* small RNA sequencing datasets and data from *B. rapa* anthers we defined 84,468 small RNA loci covering a total of 102.5 megabases (29.6%) of the *B. rapa* R-o-18 genome. These loci were categorized by the most common size of small RNA reads mapping to them, and over 88% of small RNA loci contained primarily 24-nt siRNA (24-dominant) (Supplemental Figure 1).

We began by comparing expression in cells from non-fertilized organs: ovules, which are composed almost entirely of diploid integument cells, and leaves. We examined the distribution of small RNA accumulation at expressed loci in these two tissues and found that, despite a larger number of loci with detectable small RNA accumulation in leaves, the highest RPKM values were from ovules (Figure 1A). This observation suggests that a few highly expressed loci might be obscuring representation of other loci in our ovule siRNA libraries.

**Figure 1.**
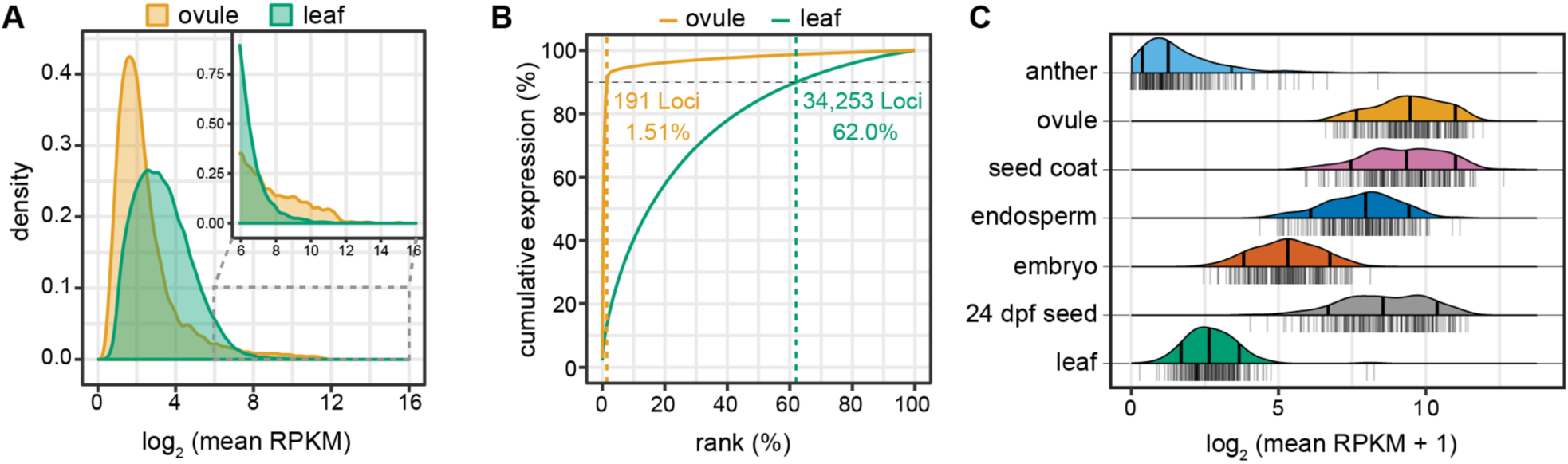
Siren loci dominate the small RNA expression landscape in seeds. (A) Distribution of small RNA accumulation at each locus in ovule and leaf. The average of expression for loci with at least 2 replicates ≥ 2 RPKM is plotted (n = 11,149 in ovules, 61,721 in leaves). Despite the higher median expression in leaves, the highest expressed loci are found in ovules (inset). (B) Cumulative expression plot of loci in ovule and leaf. Total expression at each 24-dominant locus was determined in reads per million (RPM) using combined replicates, ranked, and their cumulative expression was plotted. Loci were analyzed if they had expression ≥ 2RPM in combined replicates (n = 12,636 in ovules, 55,228 in leaves). The 1.51% highest expressed 24-dominant loci are responsible for 90% of small RNA expression in ovules. In leaves, the highest 62% of 24-dominant loci account for 90% of expression. (C) Mean small RNA accumulation at the 191 siren loci in several tissues. Average expression level from replicates is shown, with individual data points plotted as a rug underneath the ridge plot. Quantile lines show 10th, 50th (median), and 90th percentiles.

To identify which loci are responsible for the largest share of small RNA abundance in ovules we compared the cumulative expression distribution for 24-dominant loci in ovules versus leaves (Figure 1B). Comparing the contribution of each 24-dominant locus to the cumulative expression in leaf and ovule reveals a stark difference between the organs. Ninety percent of the expression in ovules was derived from only 191 loci (1.51% of ovule-expressed loci). In comparison, 34,424 loci (62.0%) are required to reach 90% of cumulative expression in leaves.

We next compared small RNA abundance at these highly expressed ovule loci in multiple organs (Figure 1C). The 191 loci are highest expressed in ovule and seed coat. Ovules are predominantly composed of integument, which is the precursor of the seed coat, suggesting that expression of siren siRNAs begins in this maternal sporophytic tissue before fertilization, and continues after fertilization. SiRNAs from the 191 loci accumulate at negligible levels in leaves and anthers, demonstrating that their expression is associated with maternal reproductive structures before fertilization. However, siRNAs from the 191 loci also accumulate moderately in endosperm and to a low level in embryos, and remain expressed throughout the latest stages of embryogenesis, including in mature seeds (Figure 1C), indicating that they could affect targets throughout all stages of seed development.

The high expression of this small subset of ovule loci in endosperm indicates that they are similar to a class of small RNA loci identified in rice endosperm (Rodrigues et al., 2013), termed siren loci (<u>s</u>mall-<u>i</u>nterfering <u>R</u>NA in <u>en</u>dosperm). While our analysis indicates that siren loci are not specifically expressed in the endosperm, the previous study did not interrogate expression in ovules or seed coats and could not have observed accumulation of siren siRNAs in these tissues. We therefore refer to the 191 highly expressed ovule small RNA loci as siren loci to indicate their congruence with this previous work, and suggest that the term “siren” might better indicate a prominent siRNA signal rising above all others.

### *B. rapa* siren loci have unique sequence characteristics compared to most 24-dominant siRNA loci

To further characterize the siren loci we compared their size to other categories of small RNA loci. These included 24-dominant loci which were not classified as siren loci, 21- and 22-dominant loci, and 100 randomly sampled sets of 24-dominant loci (n = 191). The median size of siren loci was 4,238 bp, and was significantly greater than all comparison groups (Figure 2A). Siren loci account for 959.5 kb of genomic sequence (0.28%) and 0.94% of the total size of the small RNA-producing portion of the genome. However, we noted that siRNA accumulation in sirens was not uniform. Generally, most siRNAs mapped to a “core” region, with lower accumulation in flanking regions (Supplemental Figure 2). These core regions are substantially smaller (Figure 2A), but still account for 87% of siRNA accumulation at all sirens.

**Figure 2.**
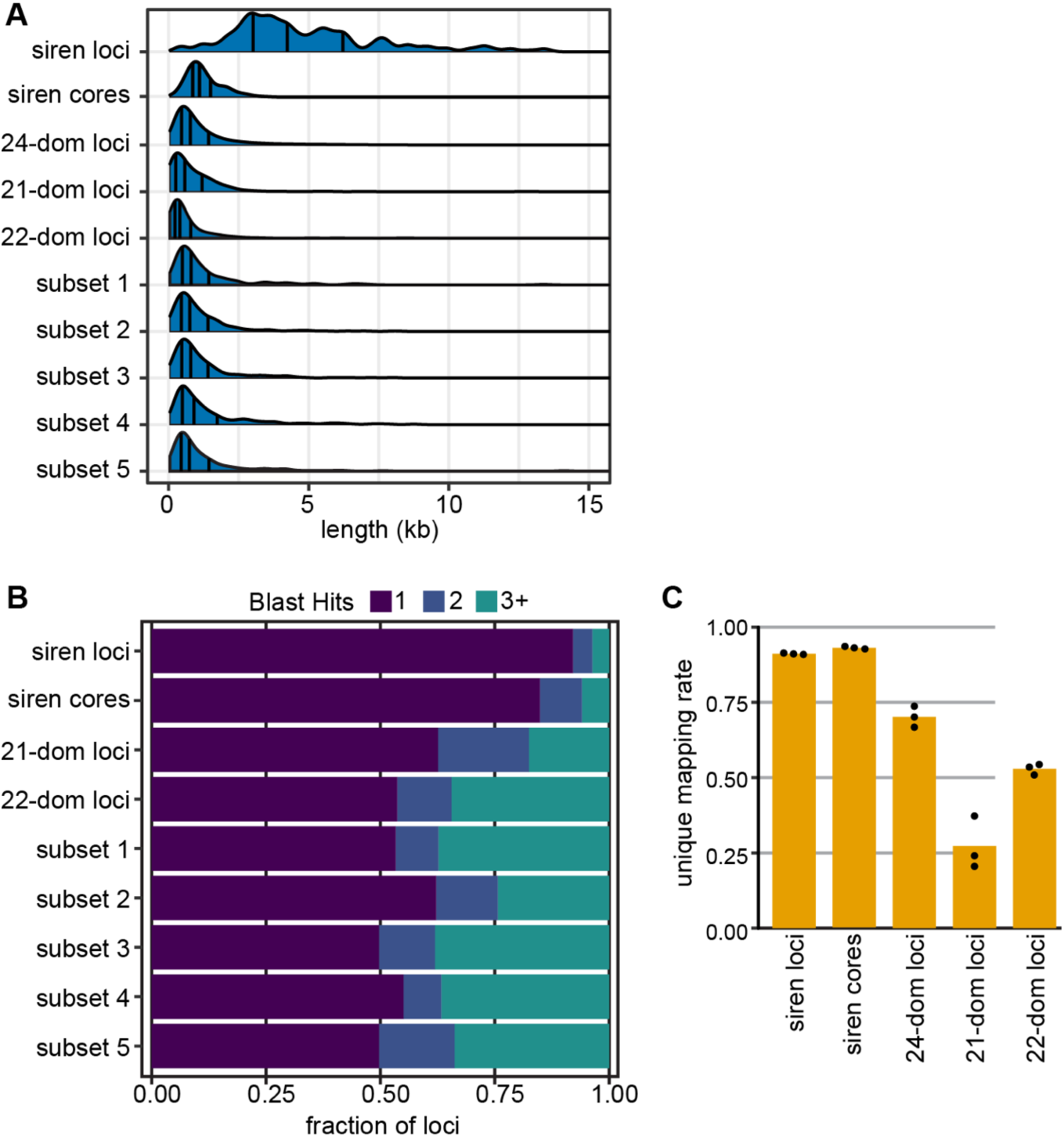
Siren loci are large and unique. (A) Length distribution of small RNA loci. Siren loci are larger than other categories, including non-siren 24-dominant (n = 74,269), 21-dominant (n = 503), 22-dominant (n = 990), and 100 random subsets of 24-dominant loci (n = 191 each, 5 shown). Quantile lines represent 10th, 50th (median), and 90th quantiles. (B) Valid blast hits by small RNA locus category. Most siren loci are present in single-copy genome-wide, as determined by blastn search against the *B. rapa* R-o-18 genome. (C) Fraction uniquely mapping reads aggregated across all loci per category, in ovule samples. Siren loci contain more uniquely mapping reads than non-siren 24-dominant, 21-dominant, and 22-dominant loci. Bars represent the mean of three replicate libraries, individual values shown.

To assess the copy number of siren loci we performed a BLAST search against the *B. rapa* genome using all 191 siren loci and the same comparison groups described above as queries. BLAST hits were collected if they were ≥ 432 nt (the first quartile for length of all *B. rapa* small RNA loci), ≥ 50% of the query locus length, had ≥ 80% identity, and an E value ≤ 1×10^−50^. With these criteria, 90% of siren loci had only a single BLAST match compared to approximately 50% of comparison groups (Figure 2B). Siren loci also had fewer blast hits than the 191 random loci of equal sizes. Consistent with the uniqueness of siren loci, the siRNAs mapping to these loci are more likely to map uniquely in the genome compared to other categories of loci (Figure 2C).

Next, we assessed overlap between small RNA-producing loci and a variety of genomic annotations (Table 1). Siren loci show neither depletion nor enrichment of genes compared with randomized coordinates of the same size, and therefore are enriched for genes relative to other 24-dominant siRNA loci, which show depletion for genes (Table 1). Siren loci are also enriched for DNA elements (class II TEs) and helitrons and depleted for retrotransposons (class I TEs). However, when only siren core sequences are considered, genes are depleted and helitrons are the only transposon class that is enriched. (Table 1). This further highlights the uniqueness of siren loci in the genome and a potential interaction, specifically, with helitron-type DNA TEs.

**Table 1.** Overlap between siRNA loci and genomic features.

|  | Genes | TEs | Class I<br>TEs | Class II<br>TEs | Helitrons |
| --- | --- | --- | --- | --- | --- |
| **24-dominant (n = 74,269)** |  |  |  |  |  |
| Overlapping loci<sup>1</sup> | 34.8% | 89.7% | 33.4% | 45.0% | 28.0% |
| Fold enrichment vs genome | **0.7** | **1.4** | **1.1** | **2** | **2.1** |
| **siren loci (n = 191)** |  |  |  |  |  |
| Overlapping loci<sup>1</sup> | 75.9% | 96.3% | 34.0% | 67.0% | 62.8% |
| Fold enrichment vs genome | 1.1 | **1.1** | **0.7** | **1.5** | **2** |
| Fold enrichment vs 24-nt dominant | **2.1** | 1.1 | 1 | **1.5** | **2.2** |
| **siren cores (n = 191)** |  |  |  |  |  |
| Overlapping loci<sup>1</sup> | 39.8% | 70.7% | 7.85% | 27.7% | 40.8% |
| Fold enrichment vs genome | **0.8** | 1.1 | **0.3** | 1.2 | **2.8** |
| Fold enrichment vs 24-nt dominant | 1.1 | **0.8** | **0.2** | **0.6** | **1.4** |

<sup>1</sup> overlap of $\geq 1$ nucleotide required **bold**, $p < 0.0001$

Because we categorized loci as 24-dominant when a plurality of their siRNAs were 24-nt, we also compared the fraction of RNAs between 19 and 26 nt in length that accumulated at siren loci (Figure 3A). Both siren and non-siren 24-dominant loci have a high proportion of 24-nt RNAs, whether assessed in aggregate (Figure 3A) or individually (Figure 3B).

**Figure 3.**
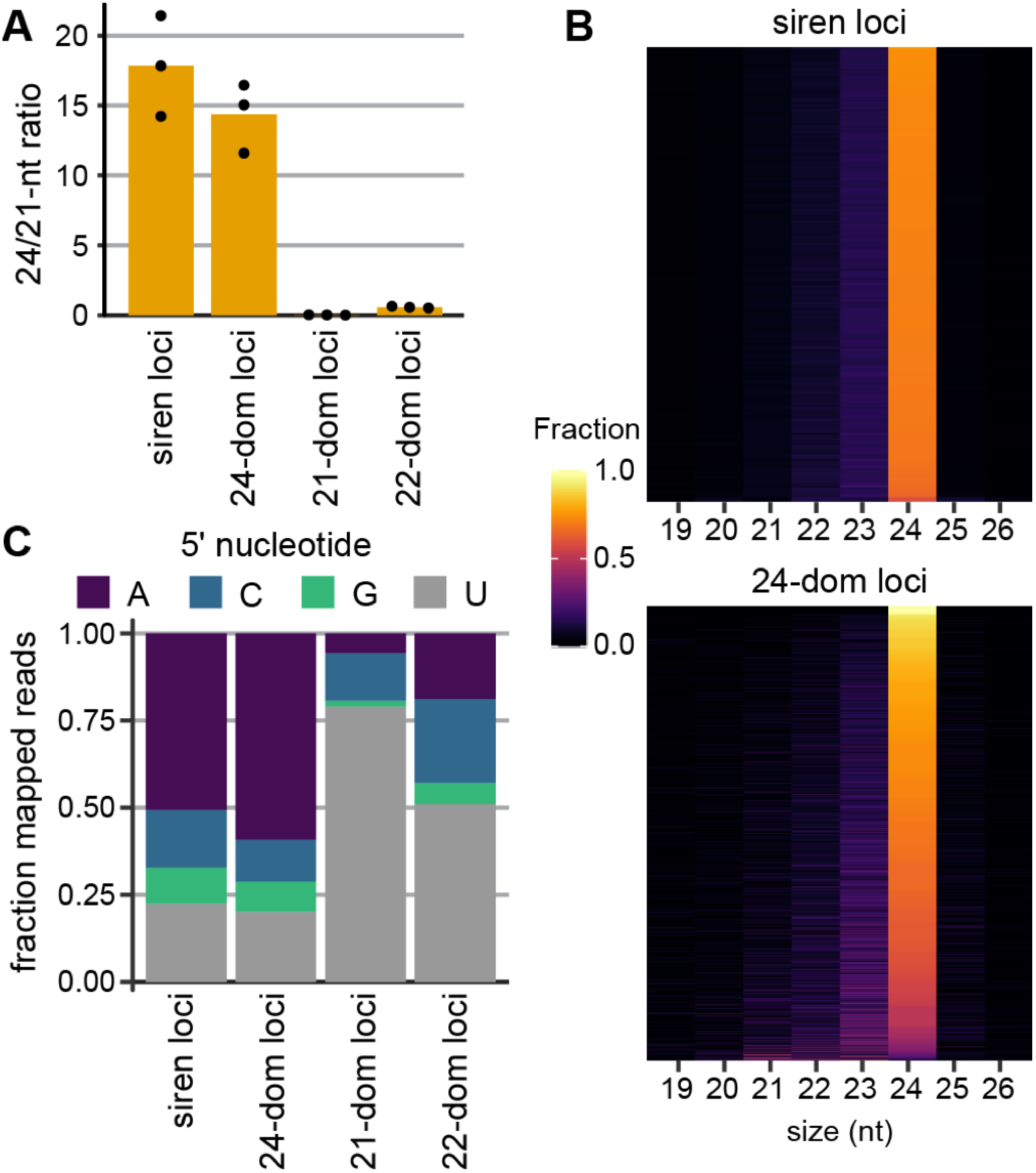
Siren loci accumulate 24-nt small RNAs beginning with adenine. (A) 24/21-nt ratio of small RNAs, aggregated across all loci of a category. Bars represent the mean of three ovule libraries, individual values shown. (B) Heatmap showing the fraction of each size class of RNA present at each locus (y-axis). (C) 5’ nucleotide bias of all RNAs in each category. (B, C) are from from combined ovule libraries.

Small RNAs function when bound to Argonaute proteins, which have distinct 5’ nucleotide preferences. The primary Argonaute associated with 24-nt siRNAs and RdDM is AGO4, which exhibits an adenine bias at the 5’ nucleotide (Frank et al., 2012; Mi et al., 2008). Siren siRNAs have the same 5’ adenine bias as siRNAs from 24-dominant loci, suggesting that siren RNAs may interact with AGO4 or related Argonaute proteins (Figure 3C).

The size of siren loci, their single-copy nature, and differences in associated genomic features together argue that siren loci are distinct from the majority of 24-dominant loci. However, accumulation of 24-nt siRNAs that begin with adenine suggests siren loci might interact with AGO4 and target *de novo* DNA methylation (Havecker et al., 2010; Mi et al., 2008).

### Siren siRNAs direct DNA methylation

To determine the biogenesis of siren siRNAs we compared small RNA accumulation at siren loci in *B. rapa nrpd1, rdr2*, and *nrpe1* mutants. To address oversampling that results from loss of most siRNAs in *nrpd1* and *rdr2* mutants, small RNA accumulation was normalized to mapped 21-nt RNAs, which are not affected by mutations in the RdDM machinery (Herr et al., 2005; Kasschau et al., 2007). Median siren siRNA accumulation was reduced by approximately 16-fold in *nrpd1* ovules, and 8-fold in *rdr2* ovules, while *nrpe1* ovules maintain wild-type expression of siren RNA (Figure 4A). This pattern of expression indicates that siren siRNAs are canonical Pol IV/RDR2 products that are not dependent on Pol V (Mosher et al., 2008).

**Figure 4.**
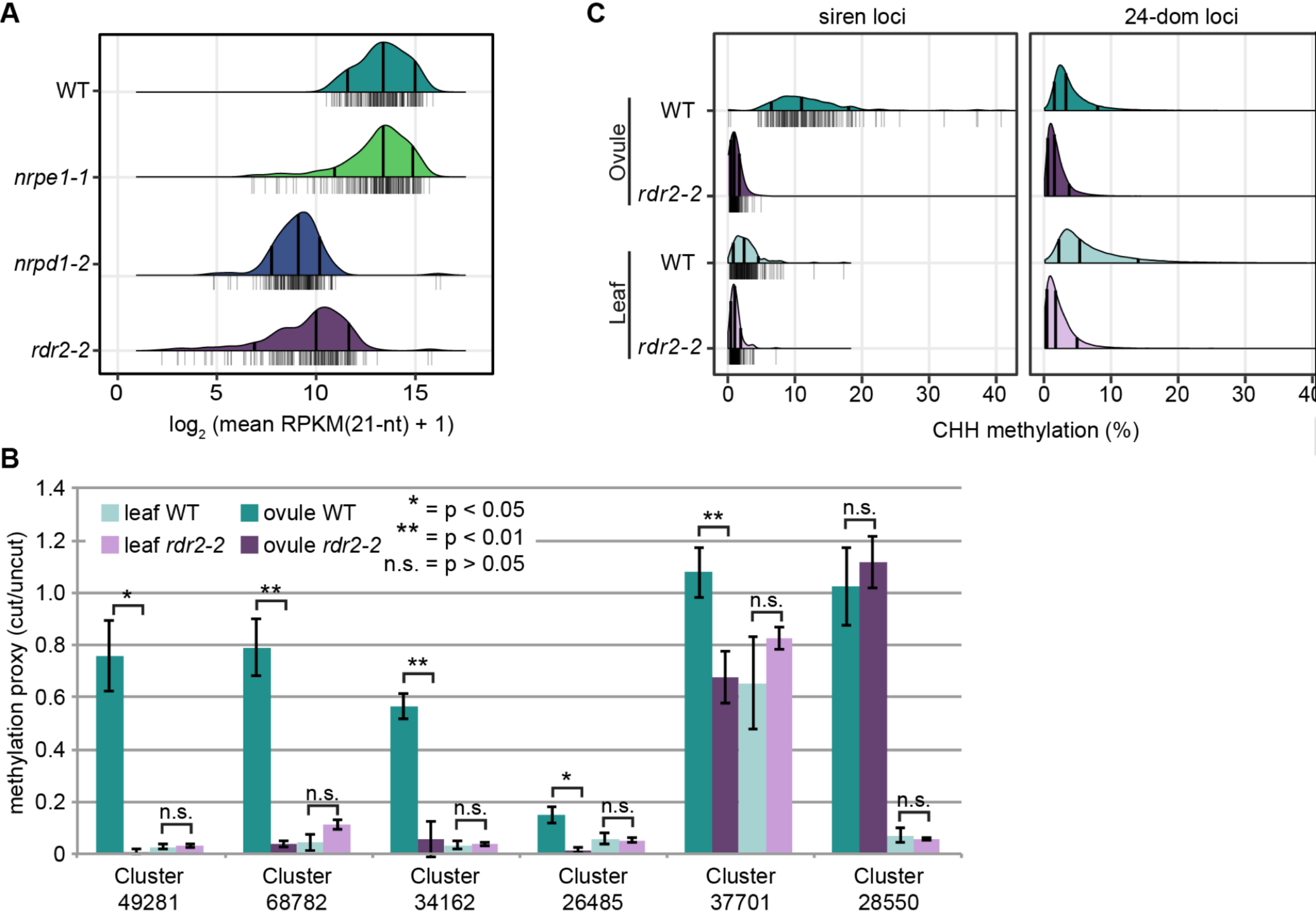
Siren loci are organ-specific RdDM targets. (A) Small RNA accumulation at siren loci in ovules. Small RNA abundance at each locus is normalized by locus length and million mapped 21-nt reads per library. Mean of three replicates shown. Quantile lines show 10th, 50th (median), and 90th percentiles. Individual measurements shown as the rug below each density. (B) Relative methylation from chop-qPCR. Bar is the mean of three replicates, error bars represent standard deviation. Five of six tested siren loci are significantly differentially methylated at the assayed nucleotide based on a two-tailed Student’s t-test. (C) CHH methylation at siren loci and non-siren 24-dominant loci. Percent methylation calculated based on methylated and unmethylated cytosine calls over each locus in the category and sample. Minimum depth of 5 reads required to calculate a percentage.

To determine, unequivocally, whether siren loci are associated with RdDM, we first assessed DNA methylation at six randomly-selected siren loci using methylation-sensitive quantitative PCR (Figure 4B). This method assesses methylation of individual cytosines in the asymmetric CHH context (where H is A, T, or C) and the results are therefore indicative of *de novo* methylation (Law and Jacobsen, 2010; Zhang et al., 2014). The level of DNA methylation at these sites is lower in leaves compared to ovules, correlating with expression of siren siRNAs (Figure 2A). At 5 of 6 loci tested, *rdr2* shows a significant reduction in DNA methylation in ovules, consistent with our expectation that siRNA accumulation enables DNA methylation at siren loci.

To investigate DNA methylation genome-wide, and at all siren loci, we performed whole-genome bisulfite sequencing in wild-type and *rdr2* ovules and leaves. Consistent with the methylation-sensitive PCR assay, siren loci exhibit more CHH methylation in ovules than leaves and this methylation is lost in *rdr2* ovules (Figure 4C). Compared to non-siren 24-dominant loci, siren loci are highly methylated in ovules, which might be a consequence of the substantial accumulation of siRNAs at these loci. This result, in combination with the small RNA sequencing, establishes siren loci as organ-specific sites of RdDM.

### Siren loci are present across angiosperms

To determine whether siren loci are specific to *B. rapa*, or are present in more diverse species, we utilized public data from *A. thaliana* and rice (Autran et al., 2011; Baldrich et al., 2019; Huang et al., 2019; Kirkbride et al., 2019; Rodrigues et al., 2013). Using these datasets we identified 16,422 small RNA loci in *A. thaliana* and 122,661 loci in rice. We then compared the contribution of each locus to the overall small RNA accumulation in that tissue as we previously did in *B. rapa*.

Similar to *B. rapa, A. thaliana* ovules were dominated by accumulation of siRNAs at a small number of loci (Figure 5A). Similar profiles for the same tissue from different public datasets demonstrate that cumulative expression patterns are robust to factors such as library prep kit, sequencing instrument, and which lab performed the experiment. Using the same 90% cumulative ovule expression threshold as in *B. rapa*, we identified 128 ovule siren loci in *A. thaliana* and assessed accumulation of siRNAs at each locus. As in *B. rapa*, accumulation at these loci remained high in developing seeds, but was substantially reduced in leaves and in *nrpd1* and *rdr2* seeds (Figure 5B, C). Combined with the expression bias in ovules, this genetic and developmental expression pattern confirms that these are siren loci.

**Figure 5.**
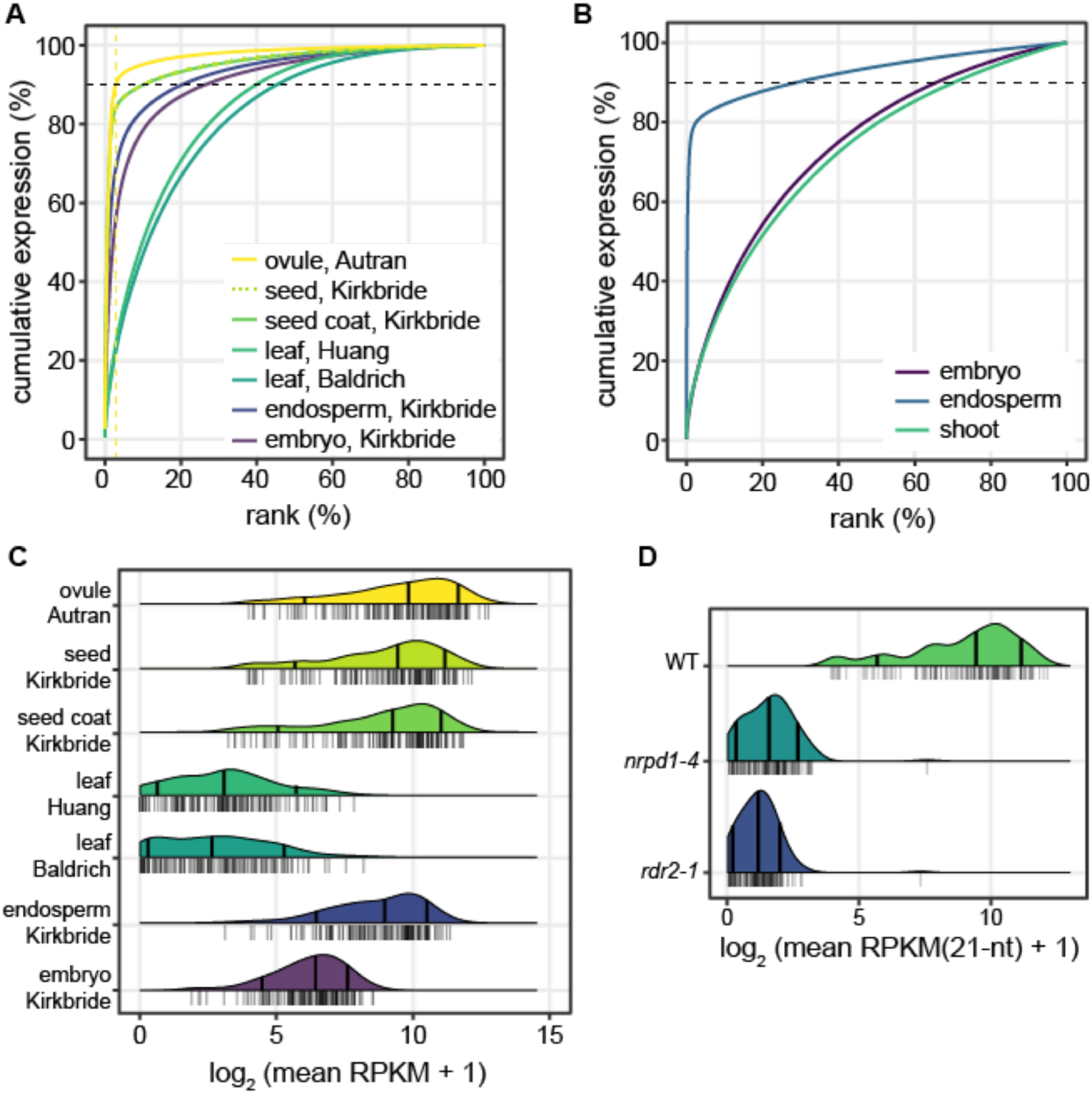
Siren loci are present in diverse angiosperms. (A) Cumulative expression plot as in Figure 1A in publicly available *A. thaliana* datasets. *A. thaliana* siren loci are defined by 90% of cumulative expression in the ovule dataset. (B) Cumulative expression plot for *O. sativa* small RNA data. (C) Ridge plot showing mean small RNA accumulation at *A. thaliana* siren loci in several tissues. (D) Ridge plot showing mean small RNA accumulation at *A. thaliana* siren loci in seeds. Samples taken from the Kirkbride *et al.* 2019 dataset. For all ridge plots, quantile lines show 10th, 50th (median), and 90th percentiles. Individual data points are the mean of available replicates, which are plotted as a rug underneath each ridge plot.

In rice endosperm we identified a similarly skewed cumulative expression pattern (Figure 5D), as we expected due to the previous observation of biased siRNA accumulation in this tissue (Rodrigues et al., 2013). The unequal accumulation of siRNA in diverse plants indicates that extreme expression of siRNAs from a few siren loci is a conserved feature of angiosperm seed development.

### Siren loci exhibit high rates of evolution

To investigate the evolutionary rate of siren loci, we searched for homology between *B. rapa* and *A. thaliana* siren loci. Because *B. rapa* experienced a recent whole genome triplication approximately 22 million years ago, *A. thaliana* loci are homologous with up to three loci in the *B. rapa* genome (Beilstein et al., 2010; Wang et al., 2011). Using homology of protein-coding genes as a guide, syntenic location(s) in the *B. rapa* genome were identified for the 64 highest-expressed *A. thaliana* sirens. None of these locations showed sequence similarity to the *A. thaliana* siren loci. However, 9 (14%) had a siren locus at one or more *B. rapa* syntenic positions (Supplemental Table 1). One *A. thaliana* siren was conserved at all three *B. rapa* orthologous positions, despite lack of nucleotide sequence conservation among these positions (Figure 6). Seven additional *B. rapa* siren loci were conserved at a homeologous position in *B. rapa*, and, with the exception of one pair, these also had no conserved sequence. These data demonstrate that over the 40 million years since the divergence of *A. thaliana* and *B. rapa* from a common ancestor (Beilstein et al., 2010), a siren’s position is only rarely conserved and sequence is not conserved. In addition, comparison of homeologous sequences within *B. rapa* indicates that nucleotide sequence at siren loci has diverged rapidly in the approximately 22 million years since the whole genome triplication.

**Figure 6.**
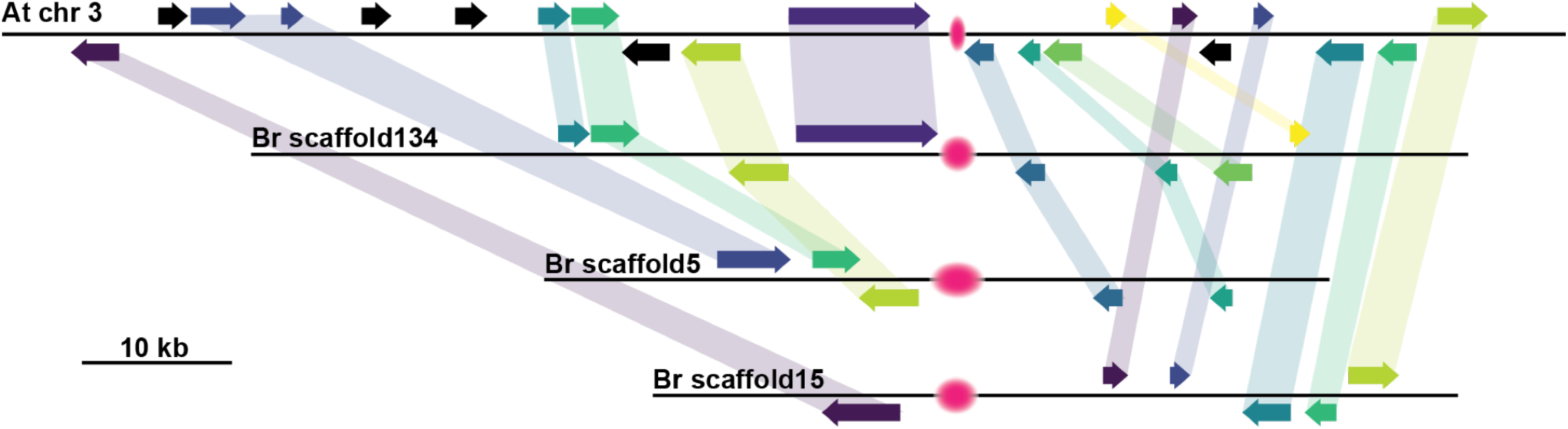
Siren loci can be conserved between *B. rapa* and *A. thaliana*. (A) Depiction of synteny between three homeologous regions of the *B. rapa* genome and a homologous region of *A. thaliana* chromosome 3. All three homeologous regions in *B. rapa* and the orthologous region of the *A. thaliana* genome contain a siren locus, depicted as a pink fuzzy oval. Homologous genes are shown with identically colored arrows and shadows.

### Siren RNAs are maternal in origin

The primary function of RdDM is hypothesized to be silencing of transposons (Matzke and Mosher, 2014). Because the seed coat is destined for programmed cell death, and therefore control of transposons is not necessary, it is counter-intuitive that siren siRNAs would express highly in this tissue. Alternatively, siRNAs might be produced within the seed coat in order to transit into the seed where they are required for development of embryo or endosperm. If siren siRNAs that accumulate in filial tissues are produced in the maternal sporophyte, they should map specifically to maternal chromosomes.

To determine the parental origin of siren RNAs, we sequenced small RNAs from reciprocal crosses between two *B. rapa* varieties, R-o-18 and R500. We then determined the parental contribution of siRNAs arising from the R-o-18 siren loci and their R500 counterparts by counting reads that map specifically to only one variety and fall within a siren locus. In R-o-18 × R500 endosperm, approximately 4.5% of siren RNAs mapped specifically to R-o-18, a level similar to R-o-18 × R-o-18 endosperm and significantly higher than R500 × R-o-18 endosperm (Figure 7A). Similarly, approximately 2.3% of siren siRNAs in R500 × R-o-18 endosperm were R500-specific, a fraction that was unchanged when R-o-18 genomes were contributed from the pollen. Siren siRNAs are strongly maternal in hybrid endosperms, as the paternal genotype does not influence the percentage of maternal-specific siren siRNAs.

**Figure 7.**
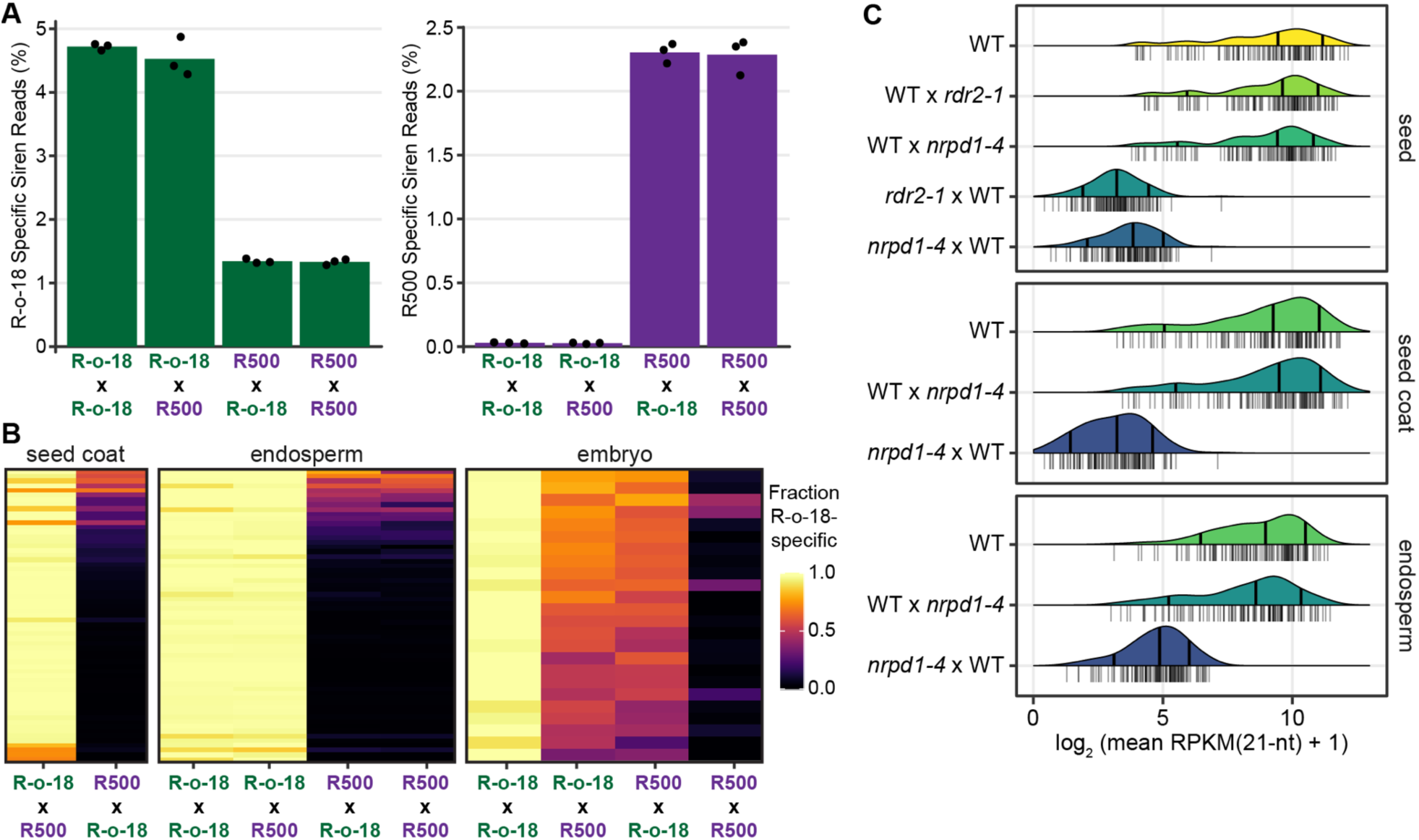
Siren RNAs are maternal in origin. (A) Percentage R-o-18-specific and R500-specific siren reads in endosperm of R-o-18 × R500 reciprocal crosses. Bars represent mean of three replicates, individual replicates are plotted as dots. The 1% of R-o-18-specific siren reads in the R500 × R500 cross likely derive from regions of the R500 genome that produce siRNAs but are not present in the current R500 genome assembly. (B) Heatmap showing the fraction of R-o-18-specific reads at each *B. rapa* siren locus in reciprocal R-o-18 × R500 crosses, in seed coat (n = 64), endosperm (n = 62), and embryo (n = 24). Loci were analyzed if they had ≥10 reads across all samples in a tissue and an R-o-18 read fraction < 0.7 in R500 × R500 (R500 × R-o-18 for seed coat), as R-o-18 specific reads in those samples represent reads derived from regions also present in R500 but absent from the current version of the R500 genome assembly. (C) Ridge plot of small RNA accumulation at *A. thaliana* siren loci in endosperm, seed coat, and whole seeds. Data from Kirkbride *et al.* 2019 dataset. Quantile lines show 10th, 50th (median), and 90th percentiles. Individual data points are the mean of available replicates, plotted as a rug underneath each ridge plot.

We further assessed the parental origin of siren siRNAs in the endosperm by measuring maternal bias at individual siren loci (Figure 7B). Maternal bias in the endosperm was indistinguishable from the maternal sporophytic seed coat, suggesting that siren loci are either expressed only from matrigenic (maternally-inherited) alleles in the endosperm or that siRNAs produced in the developing seed coat accumulate in the endosperm. In contrast to the marked maternal bias in endosperm, siRNAs were biallelically expressed in embryos (Figure 7B), indicating that siren siRNAs are expressed from both matrigenic and patrigenic alleles in the embryo.

To eliminate the possibility that maternal bias of siren siRNAs in the endosperm was due to contamination by maternal seed coat (Schon and Nodine, 2017), we measured siren accumulation in publicly-available *A. thaliana* libraries, including laser-capture microdissected endosperm (Kirkbride et al., 2019). Reciprocal crosses between *nrpd1* or *rdr2* and wild type demonstrate that siRNA accumulation at siren loci in whole seeds or seed coats is dependent on maternal alleles of *NRPD1* and *RDR2* (Figure 7C). In endosperm, siRNA accumulation at siren loci was dependent only on maternal *NRPD1*.

Together, these results suggest that, although siren loci express siRNA from both matrigenic and patrigenic alleles in the embryo, endosperm accumulates only maternal siren siRNAs. Because the majority of the siRNA accumulation in developing seeds is from siren loci in endosperm and seed coat, developing seeds are therefore dominated by maternal-specific siRNAs.

## Discussion

Here we have shown that siRNA accumulation in seeds is overwhelmingly due to siren loci, a small number of highly-expressed siRNA loci. Siren siRNAs are most abundant in ovules before fertilization and comprise 90% of siRNA accumulation in that tissue. Strong expression of siren loci in maternal sporophytic tissue after fertilization explains conflicting reports regarding maternal siRNA bias in whole seeds (Mosher et al., 2009; Pignatta et al., 2014), as these sporophytic siRNAs mask the maternal bias from the same loci in endosperm.

An alternative explanation for the abundant siren siRNAs in ovules is that all other siRNA loci are downregulated, leaving only siren loci remaining. Two observations argue against this hypothesis. Firstly, abundant 24-nt siRNA accumulation is routinely detected in reproductive tissues from a variety of plants, indicating that widespread downregulation of 24-dominant loci is not a general phenomenon (Hu et al., 2015; Huang et al., 2013; Liu et al., 2014; Nystedt et al., 2013). Secondly, siren siRNAs are abundant in *B. rapa* whole seeds at the mature green stage (Figure 1C), despite the fact that these seeds are comprised mostly of embryo, which expresses a more diverse population of siRNAs. The abundance of 24-nt siRNAs in reproductive tissues and accumulation of siren siRNAs in late-stage seeds argues that skewed expression patterns in ovules and seed coats results from specific upregulation of siren loci rather than downregulation of all other loci.

Exactly what distinguishes siren loci from most RdDM loci is not clear. While other RdDM loci are enriched for all classes of transposons, siren loci are enriched only for helitrons sequences. Unlike other RdDM loci, siren loci are specifically expressed in ovules and developing seeds, and likely represent the “Type I” loci previously described as reproductive-specific in *A. thaliana* (Mosher et al., 2009). How this specific and strong expression is established is unclear, since the only known mechanism of Pol IV recruitment is via the chromatin reader SAWADEE HOMEODOMAIN HOMOLOG 1 (SHH1), which recognizes the combination of unmethylated Lysine 4 and methylated Lysine 9 on Histone H3 (Law et al., 2013). However, SHH1-mediated recruitment only accounts for about half of known RdDM loci, suggesting that additional mechanisms of Pol IV recruitment remain to be identified. Other examples of organ-specific RdDM suggest that regulated expression of Pol IV activity, perhaps through tissue-specific epigenetic modification, is an underexplored developmental mechanism (Huang et al., 2019; Walker et al., 2018).

We were surprised by our result that, occasionally, the syntenic chromosomal position of a siren in *A. thaliana* was conserved in *B. rapa*, but always in the absence of sequence conservation (Figure 6). We consider two possible explanations for this “position but not sequence conservation” result. Firstly, siren character (*i.e.*, abundant reproductive-specific siRNA production) might be determined by a conserved epigenetic signature, much like centromere position is determined by specific chromatin (Mendiburo et al., 2011). Alternatively, a siren control sequence might exist linked to, but not overlapping, the siren itself. This could account for conservation of siren character without conservation of siren sequence. If this siren control sequence has a “high birth and death” rate (Nam et al., 2004) siren character would also disappear at a high rate. These two explanations are not mutually exclusive.

Siren siRNAs are maternally biased in the endosperm (Figure 7). This bias might result from imprinted expression in the endosperm (matrigenic expression) or movement of siren siRNAs into the endosperm from surrounding maternal sporophytic tissue where siren siRNAs are abundantly expressed (maternal expression). In *A. thaliana*, Pol IV is required maternally for accumulation of siren siRNAs in filial tissues (Figure 7C). This observation supports the maternal expression model, however it remains possible that Pol IV is required during female gametophyte development to establish the epigenetic marks necessary for imprinted matrigenic expression from siren loci (Mosher, 2010). How siRNAs might move between the developing seed coat and filial tissues is currently unknown.

The developmental function of siren expression is unknown, but RdDM activity at siren loci might be required for successful seed development. In *B. rapa*, loss of maternal RdDM causes severe seed development defects and high rates of seed abortion regardless of the genotype of the filial tissues (Grover et al., 2018), suggesting that siren siRNA production in the developing seed coat might be required for seed development. However, silencing by siren RNAs is not required in *A. thaliana*, as *nrpd1* mutant mothers have only subtle developmental phenotypes (Grover et al., 2018; Kirkbride et al., 2019). Furthermore, rapid evolution of siren loci argues against a conserved role in regulation of developmental genes. Experiments to determine whether siren siRNAs impact developmental regulators, including imprinted expression in the endosperm, are needed to ascertain any developmental role of siren expression.

The maternal bias of siren siRNAs and their production in maternal sporophytic tissues is reminiscent of piwi-associated (pi)RNAs in *Drosophila melanogaster*. In *D. melanogaster*, piRNAs are produced in ovaries before fertilization and trigger additional piRNA production in the embryo following fertilization (Iwasaki et al., 2015). In the Drosophila genome, piRNA precursor transcripts arise from roughly 140 loci containing clusters of transposon fragments, and the resulting piRNAs silence transposons (Hirakata and Siomi, 2016; Iwasaki et al., 2015). Any transposon that is active and mobile in the genome may be “trapped” by a piRNA cluster, thereby causing the production of piRNAs capable of silencing the active element. Although most siren siRNAs perfectly match only one genomic location, it is possible that siRNAs can tolerate a few mismatches at their target loci, or that spreading of siren expression to neighboring transposons is a mechanism to produce siRNAs capable of suppressing mobile elements during embryogenesis.

The production of abundant 24-nt siRNAs from the integuments surrounding the maternal gametophyte is also reminiscent of phased 24-nt siRNA production from the tapetum surrounding the male gametophyte (Ono et al., 2018). Phased siRNAs do not require Pol IV, and are instead produced following cleavage of a long Pol II transcript. The biological role of these phased siRNAs is unknown, but it is striking that they are produced from sporophytic tissue surrounding the developing gamete, the same pattern we see from siren siRNAs. We detect elevated phasing scores at siren loci; however highly-expressed siRNA loci can be misidentified by common phasing algorithms (Polydore et al., 2018). Because Pol IV produces transcripts only slightly longer than a single siRNA (Blevins et al., 2015; Zhai et al., 2015), it is unlikely that 24-nt siren RNAs are phased. Despite this difference, siren siRNAs might represent a maternal counterpoint to paternal phased 24-nt siRNAs or paternal gametophyte-derived 21/22-nt easiRNAs (Martinez et al., 2018).

Siren loci are a conserved class of maternal, reproductive, small RNA loci that are distinguished by their tissue-specific expression. The existence of siren loci provides new context to understand uniparentally-expressed siRNAs and dynamic remodeling of the epigenome within the seed. The biological function of siren loci is currently unknown, however the correlation between siren expression and seed development in *B. rapa* opens new avenues to investigate the role of tissue-specific RdDM in reproduction and to enhance yield of grain crops.

## Methods

### Plant material and growth conditions

*B. rapa* plants were grown in a greenhouse at 18°C with at least 16 hours of light. Developing seeds were manually dissected at torpedo stage (17-20 days post-fertilization) for the collection of embryo, endosperm, and seed coat samples. Ovules and anthers were collected by dissecting pistils prior to anthesis. *B. rapa* RdDM mutants were genotyped as described previously (Grover et al., 2018). RdDM mutant lines *braA.nrpd1.a-2, braA.rdr2.a-2*, and *braA.nrpe1.a-1* are referred to as *nrpd1-2, rdr2-2*, and *nrpe1-1* respectively.

### Small RNA sequencing and analysis

Small RNA sequencing libraries were prepared using the NEBNext Small RNA kit (NEB, E7300L) according to manufacturer’s recommendations and sequenced by either the University of Missouri DNA Core or the University of Arizona Genetics Core with either an Illumina HiSeq2500 or NextSeq500, depending on the sample (Supplemental Table 2-3).

Small RNA reads were obtained from the sequencing facility and quality checked with FastQC (Anders, 2010). Adapters were removed using Trim Galore (Krueger, 2012) when necessary, with options -- length 10, and --quality 20. Following trimming, structural and noncoding RNAs, and reads mapping to the *A. thaliana* chloroplast or mitochondria genomes, were removed by aligning to the rfam database v14 (Kalvari et al., 2018a, 2018b), excluding miRNAs and miRNA precursors, and the TAIR10 (Berardini et al., 2015) chloroplast and mitochondrial genomes using Bowtie (Langmead et al., 2009). Following these steps, reads shorter than 19 nt and longer than 26 nt were removed with a Python script (fastq_length_filter.py). The remaining reads were aligned to the *B. rapa* R-o-18 genome (v2, A. Baten and G. J. King, unpublished) with Bowtie and genome-mapping reads were retained. Options for all Bowtie steps were -v 0, -m 50, --best, -a, and --nomaqround, --norc was used for rfam filtering only. Detailed information on all new data, as well as publicly-available data used in this study, are available in Supplemental Tables 2-4.

Small RNA loci were annotated with ShortStack (Axtell, 2013; Johnson et al., 2016) using filtered reads from all libraries. Options used for ShortStack were --mismatches 0, --mmap u, --mincov 0.5 rpm, and -- pad 75. Replicates were checked for consistency by principal component analysis (Supplemental Figure 3) on vst-transformed counts using R and the DESeq2 package (Love et al., 2014), and by comparison of the size profiles of mapped small RNAs between replicates. A SnakeMake workflow is available for the small RNA analysis pipeline (Bose and Grover, 2019).

Expression comparisons between wild-type samples were conducted by converting small RNA counts in each locus, per sample, into Reads Per Kilobase Mapped (RPKM) using the number of genome-mapping, filtered, reads and the size of the ShortStack predicted locus. Comparisons between wild type and mutants were normalized per million mapped 21-nt reads, rather than all reads, to address oversampling of non-24-nt reads in RdDM mutants.

Siren cores were identified by repeating ShortStack’s clustering step using --pad 1 and --mincov 2rpm with only the wild-type R-o-18 ovule small RNA alignments. The resulting small RNA loci were overlapped with identified siren loci using bedtools intersect, and the overlapping locus which accounted for the largest share of each siren’s expression was taken as that siren’s core region.

Enrichment for genomic features within small RNA loci was determined using BEDTools intersect (Quinlan and Hall, 2010). Features were considered to overlap a locus if there was at least 1 nt shared. Shuffling of locus coordinates was performed using BEDTools shuffle, and random 24mer loci were selected using R software. Fold enrichment was calculated compared to the intersecting features from the random sets of loci, and z-scores were calculated, where Z = (observed - mean_random_) / standard deviation_random_. Z-scores were converted to p-values using R.

Parent-specific small RNAs were determined by first aligning small RNA reads to a concatenated R-o-18 + R500 genome (v1.4, K Greenham and R McClung, unpublished) with bowtie to calculate the number of all mapped reads. Then, uniquely-aligning (and hence, parent-specific) reads were remapped with ShortStack. ShortStack’s --locifile option was used to tabulate small RNA counts at previously defined small RNA loci in each sample.

We identified homologous siren loci in the R500 genome by BLAST search, retaining the longest BLAST hit with the highest percent identity. Using this strategy we were able to identify a homologous R500 siren locus for all 191 R-o-18 siren loci.

### Whole-genome bisulfite sequencing and analysis

Genomic DNA was extracted with the GeneJET Plant Genomic DNA Purification Kit (Thermo Fisher Scientific, K0791) and whole-genome bisulfite libraries were prepared as previously described (Urich et al., 2015). Unmethylated lambda phage DNA (Promega, D1521) was included in the library as a bisulfite conversion control. Paired end libraries were sequenced at The University of Arizona Genetics Core on an Illumina NextSeq500.

Reads were checked for quality with FastQC and trimmed with Trim Galore using options --trim-n and -- quality 20. Trimmed reads were aligned to the *Brassica rapa* R-o-18 genome (v2, A Baten and G.J. King, unpublished) with bwa-meth (Pedersen et al., 2014) Picard Tools (Broad Institute, 2019) was used to mark PCR duplicates and the properly paired alignment rate was determined with samtools (Li et al., 2009) using options -q 10, -c, -F 3840, -f 66. Genomic coverage was determined using mosdepth (Pedersen and Quinlan, 2018) with options -x and -Q 10, and a custom Python script (bed_coverage_to_x_coverage.py). Per-cytosine methylation was extracted with MethylDackel (Ryan, 2015) in two passes, first to determine the inclusion bounds based on methylation bias per read position using MethylDackel mbias, and then with MethylDackel extract. MethylDackel options were -- CHG, --CHH, and --methylKit. Bisulfite conversion rates (Supplemental Table 5) were determined by alignment to the bacteriophage lambda (NCBI Genbank accession J02459.1) and *B. rapa* var. *pekinesis* chloroplast (NCBI Genbank accession NC_015139.1) genomes and a custom Python script (bedgraph_bisulfite_conv_calc.py). All conversion rates were greater than 99%. Methylation over siren loci was determined with BEDTools intersect and a custom Python script (bedgraph_methylation_by_bed.py). Replicates were checked for consistency by plotting aggregate genome-wide methylation levels in each sequence context (Supplemental Figure 4).

A snakemake workflow for the bisulfite sequencing analysis pipeline is available (Grover, 2019).

### Methylation sensitive quantitative PCR

Genomic DNA was extracted from *B. rapa* ovules and leaves with the GeneJET Plant Genomic DNA Purification Kit (Thermo Fisher Scientific, K0791). DNA quality and concentration were determined on a Nanodrop instrument before diluting samples to equal concentrations. For each locus tested, 100 ng of genomic DNA was incubated with its respective restriction enzyme. Quantitative PCR was performed with 4 µl of digest product on a Bio-Rad MyIQ2. Cycling conditions were: 10 minutes at 95°C (enzyme activation), followed by 40 cycles of 95°C for 10s, 60°C for 30s, 72°C for 30s, and data collection. Specificity of each reaction was confirmed by the presence of a single peak melting temperature on the derivative melting curve plot, and the presence of the expected size band through gel electrophoresis of PCR products. The resulting data was analyzed relative to an undigested DNA control prepared in parallel for each locus. Primers and restriction enzymes for each siren locus tested are listed in Supplemental Table 6.

### Evolutionary analysis of small RNA loci

A list of potential *B. rapa* syntenic orthologs of *A. thaliana* was made from the combined outputs of Synfind (Tang et al., 2015) and Synmap (Lyons et al., 2008) from CoGe (https://genomevolution.org/coge/). Synfind was run under standard conditions (40 gene window, minimum of 4 anchoring genes). Synmap was run using Last (-D 25, -A 4), syntenic blocks were merged with Quota Align using a 1:3 ratio of coverage depth and an overlap distance of 40. The initial list included erroneous pairs resulting from pairing with incomplete genes or from paralagous pairs.

To remove spurious orthologs, the coverage and percent nucleotide identity for each pair of coding sequences was determined using strand-specific lastz, and putative orthologs were retained if they had an overall coverage of at least 80% and percent id of at least 70%. If a homeologous arabidopsis gene exists, it is possible that the putative orthologous pair actually has a paralogous relationship. *A. thaliana*-*B. rapa* pairs were therefore removed when the lastz percentage id and coverage scores were higher for the homeologous arabidopsis gene (Thomas, 2006). *B. rapa* genes were also removed when their percentage id fell below a certain threshold relative to the percentage id of the highest scoring rapa ortholog in each group (< max - (0.1*max)). Tandems (orthologs identified within a 10 gene window on either side) were collapsed, keeping the tandem with the highest percentage id. Since a maximum of three orthologs would be expected from the triplicated *B. rapa* genome, only the three highest scoring genes were kept.

*B. rapa* orthologs were sorted to subgenome based as described previously (Cheng et al., 2013). In the few cases where two orthologs were assigned to the same subgenome, the out-of-order ortholog was removed. Out-of-order genes in a subgenome were also removed when all four genes on either side were in-order (within 30 genes of each other) but the gene was further than 60 genes away from any of these genes.

Sirens were assigned to the closest *B. rapa* or *A. thaliana* genes using bedtools closest (Quinlan and Hall, 2010). Orthologous regions with *B. rapa* and *A. thaliana* sirens were manually inspected on CoGe GEvo panels for sirens lying between the same flanking *B. rapa* - *A. thaliana* syntenic gene pair using genomes annotated with sirens. In particularly TE-rich regions it was not always possible to identify one or more *B. rapa* syntenic region.

## Data availability

Sequencing data generated in this study was deposited in the NCBI SRA under BioProject accession PRJNA588293. A full listing of SRA accessions by sample is available in Supplemental Tables 2-5. Custom scripts used to process data are available in a GitHub repository (https://github.com/The-Mosher-Lab/grover_sirens_2019).

## Acknowledgements

We thank Dr. Robertson McClung and Dr. Kathleen Greenham from Dartmouth University for sharing the *B. rapa* R500 genome. We also thank Dr. Robert Schmitz and his lab at The University of Georgia for technical advice regarding whole-genome bisulfite library preparation. Additionally, we thank The University of Arizona Genetics Core for sequencing the samples analyzed in this study, and Maya Bose at The University of Arizona for assistance in developing the small RNA analysis pipeline. The authors are grateful to the National Science Foundation for support through grant IOS-1546825. JWG was also supported in part by NIH grant T32-GM008659.

## Author Contributions

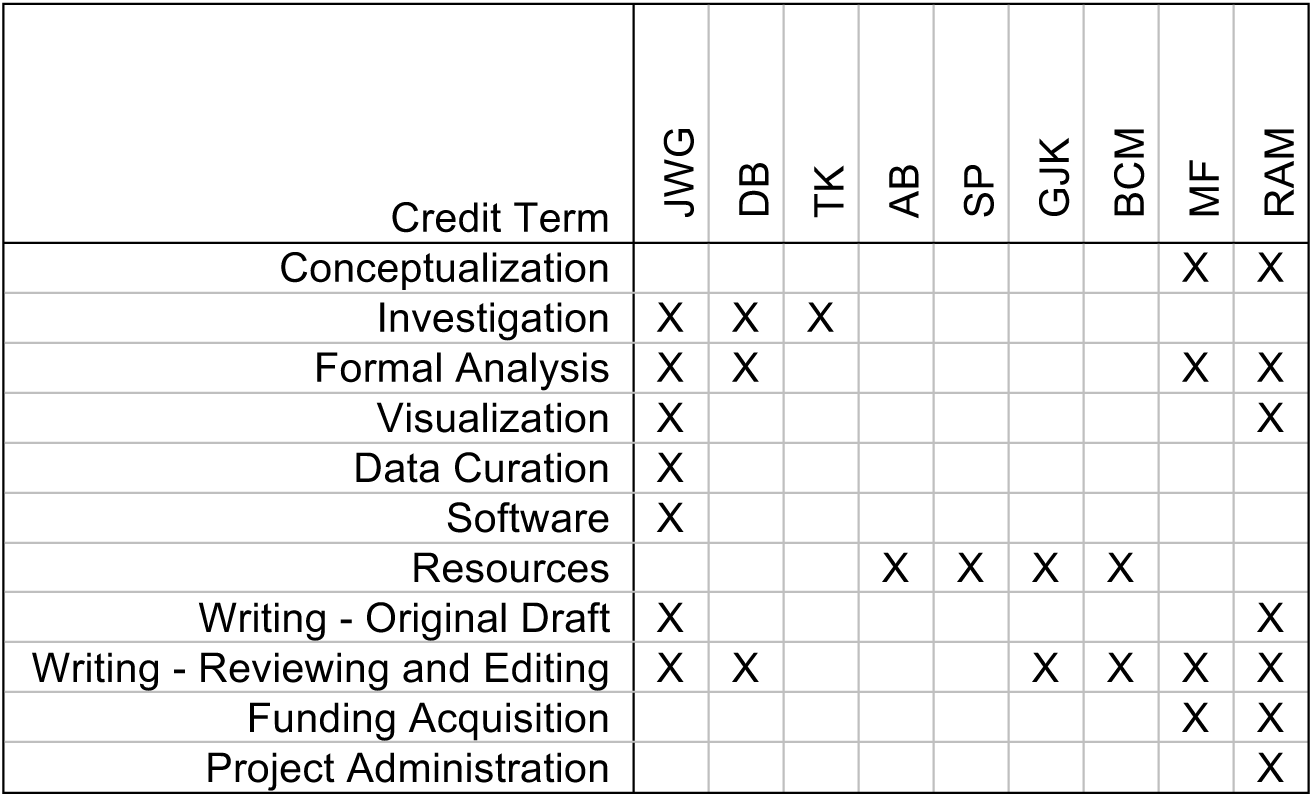

