## Supplemental Figures and Table S1 for "Abundant expression of maternal siRNAs is a conserved feature of seed development"

**Supplemental Table 1. Conservation of siren position at homeologous sites.**

| *A. thaliana* sirens | Number of *B. rapa* homeologs |
|---|---|
| 55 | 0 |
| 5 | 1 |
| 3 | 2 |
| 1 | 3 |

**Supplemental materials in additional files:**
Supplementary Table 2. R-o-18 Small RNA Sequencing Samples
Supplementary Table 3. R-o-18xR500 Reciprocal Crosses Small RNA Sequencing Samples
Supplementary Table 4. Public Small RNA Sequencing Datasets
Supplementary Table 5. Whole-genome Bisulfite Sequencing Samples
Supplementary Table 6. Methylation-sensitive qPCR primers
Data File 1: siren loci sequences

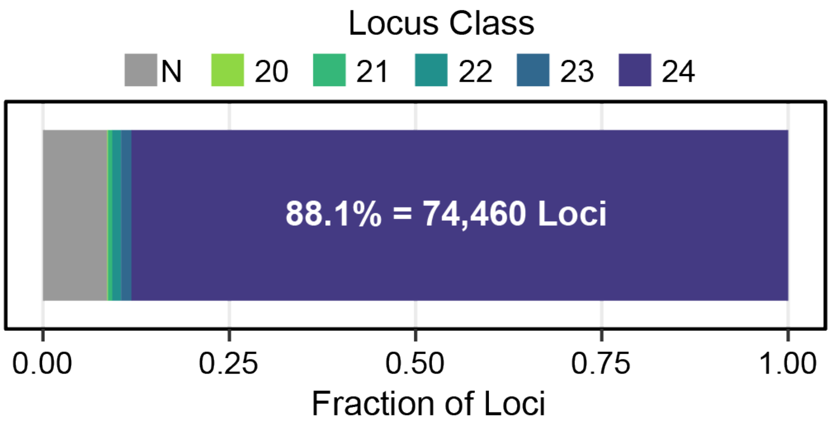


**Supplemental Figure 1. Size classification of small RNA loci in *Brassica rapa*.**
Small RNA loci generated by ShortStack are categorized by the dominant size class of RNA that accumulate over their length. We defined 84,486 small RNA loci in using our combined *B. rapa* datasets. Of these loci, 74,460 were 24-dominant, 1,173 were 23-dominant, 975 were 22-dominant, 443 were 21-dominant, and 81 were 20-dominant. A locus is given an “N” designation if 80% of the reads that accumulate over that locus are not between 20 and 24-nt in length.

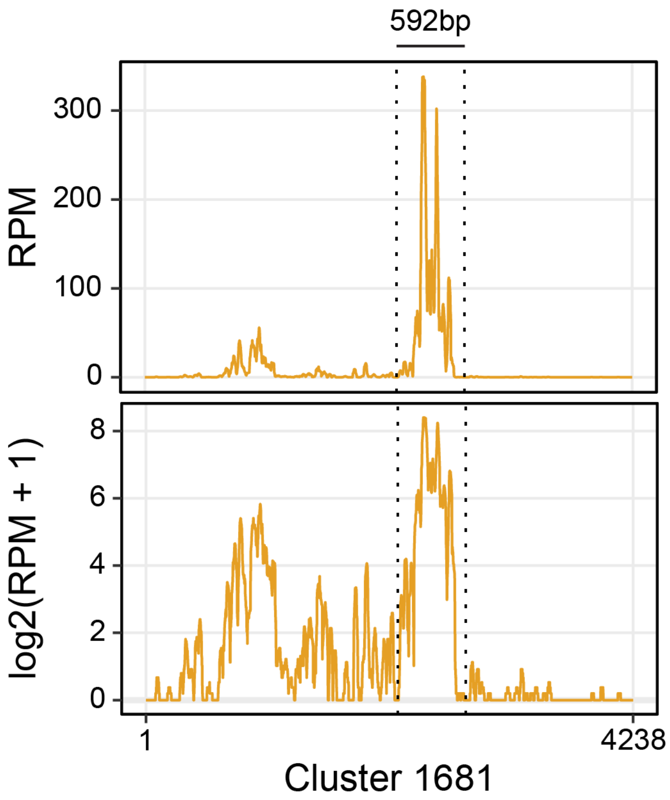


**Supplemental Figure 2. Siren cores account for most siren RNA accumulation.**
Small RNA accumulation across a single siren locus. Cluster 1681 is a siren locus with genomic coordinates scaffold1:558264-5586881. The siren core region, denoted with dotted lines, accounts for 78.5% of small RNA accumulation at this locus in ovules, and is 14% of the full siren locus length. Per-nucleotide read depth was determined with samtools depth using combined triplicate R-o-18 WT ovule small RNA alignments, and normalized to total mapped small RNAs in those samples. The same locus is shown below in log scale.

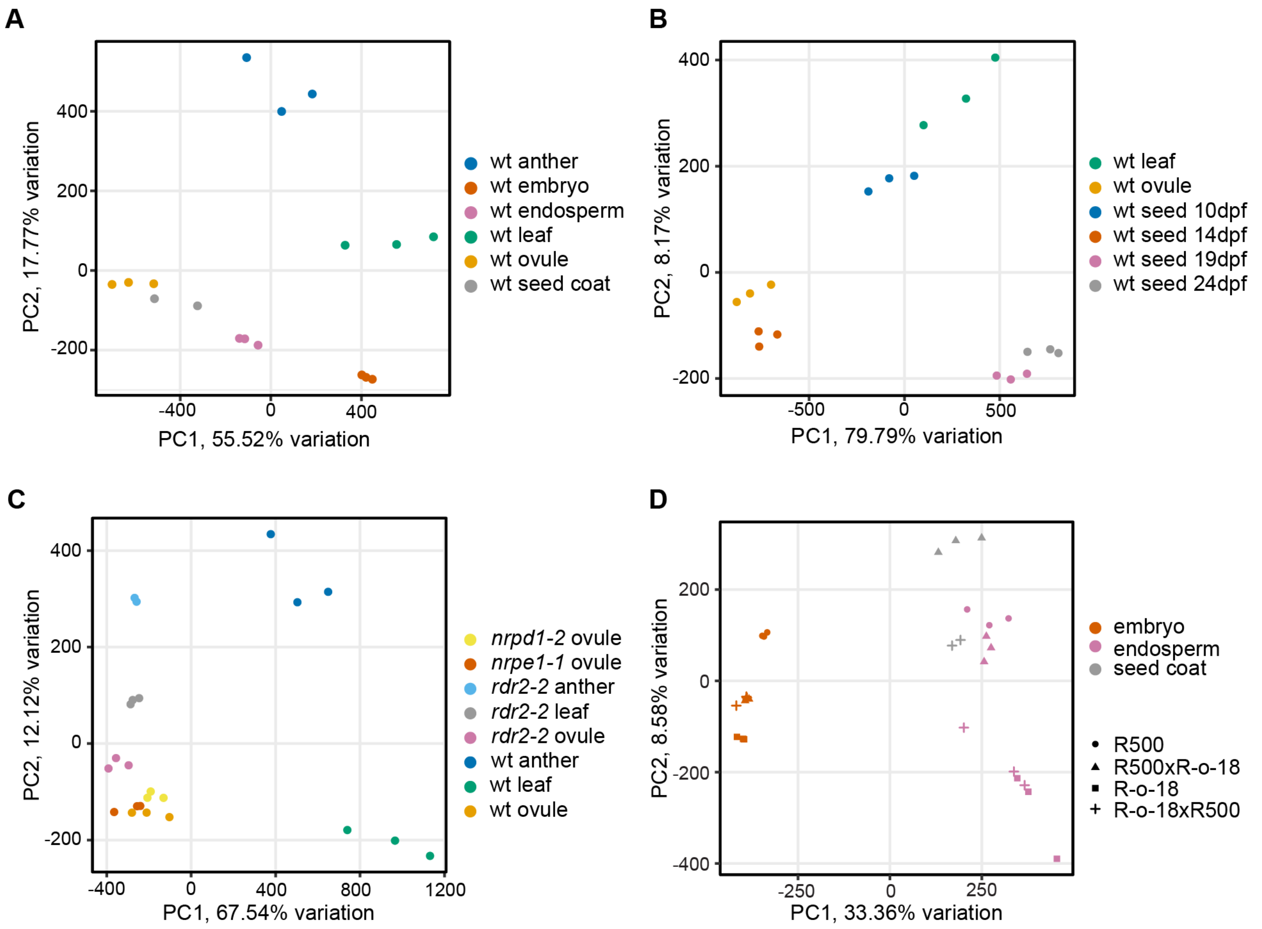


**Supplementary Figure 3. Principal component analysis of *B. rapa* small RNA samples generated in this study.**

PCA biplots from *B. rapa* small RNA samples from various organs (A), whole seed samples, ovule, and leaf (B), RdDM mutants and their corresponding wild-type controls (C), and R-o-18 x R500 reciprocal hybrids with the corresponding single-genotype controls (D). PCAs were generated from VST-transformed counts at each small RNA locus defined in *B. rapa* R-o-18 (Supplementary Figure 1). Replicates and samples which are similar primarily cluster together, showing that the replicates are consistent.

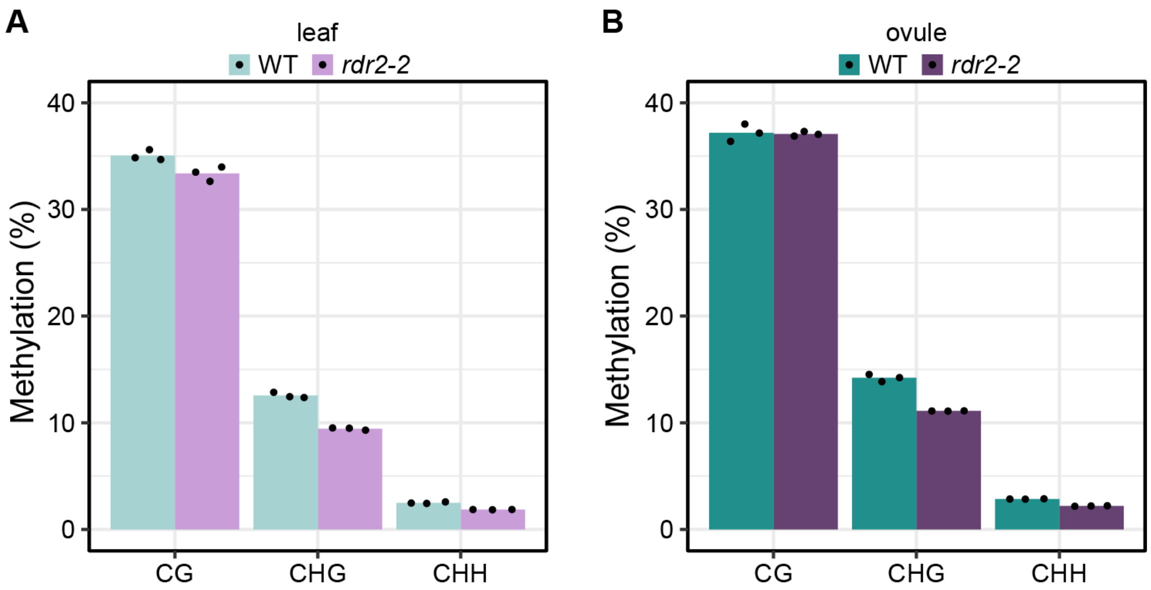


**Supplemental Figure 4. Aggregate genome-wide methylation levels in whole-genome bisulfite sequencing samples generated for this study.**

Overall methylation level in three sequence contexts in *B. rapa* wild-type and *rdr2* leaves (A) and ovules (B). Genome-wide, methylated cytosine calls were divided by total (methylated + unmethylated) cytosine calls, and expressed as a percentage. This was performed for each replicate individually. Bars represent the mean of three biological replicate libraries, individual library values are plotted as points.
